# Endocytic down-regulation of the striatal dopamine transporter by amphetamine in sensitized mice in sex-dependent manner

**DOI:** 10.1101/2023.05.17.541165

**Authors:** Tarique Bagalkot, Alexander Sorkin

**Affiliations:** Department of Cell Biology, University of Pittsburgh School of Medicine, Pittsburgh, Pennsylvania 15261

## Abstract

Dopamine transporter (DAT) controls dopamine signaling in the brain through the reuptake of synaptically released dopamine. DAT is a target of abused psychostimulants such as amphetamine (Amph). Acute Amph is proposed to cause transient DAT endocytosis which among other Amph effects on dopaminergic neurons elevates extracellular dopamine. However, the effects of repeated Amph abuse, leading to behavioral sensitization and drug addiction, on DAT traffic are unknown. Hence, we developed a 14-day Amph-sensitization protocol in knock-in mice expressing HA-epitope tagged DAT (HA-DAT) and investigated effects of Amph challenge on HA-DAT in sensitized animals. Amph challenge resulted in the highest locomotor activity on day 14 in both sexes, which was however sustained for 1 hour in male but not female mice. Strikingly, significant (by 30-60%) reduction in the amount of the HA-DAT protein in striatum was observed in response to Amph challenge of sensitized males but not females. Amph reduced Vmax of dopamine transport in striatal synaptosomes of males without changing Km values. Consistently, immunofluorescence microscopy revealed a significant increase of HA-DAT co-localization with the endosomal protein VPS35 only in males. Amph-induced HA-DAT down-regulation in the striatum of sensitized mice was blocked by chloroquine, vacuolin-1 (inhibitor of PIKfive kinase), and inhibitor of Rho-associated kinases (ROCK1/2), indicative of the involvement of endocytic trafficking in DAT down-regulation. Interestingly, HA-DAT protein down-regulation was observed in nucleus accumbens and not in dorsal striatum. We propose that Amph challenge in sensitized mice leads to ROCK-dependent endocytosis and post-endocytic traffic of DAT in a brain-region-specific and sex-dependent manner.

## INTRODUCTION

Dopamine (DA) signaling in central nervous system plays a key role in modulating cognition, locomotion, motivation, and reward-seeking behaviors (Giros and Caron, 1993; Dani and Zhou, 2004). Dysregulation of DA neurotransmission is linked to neurodegenerative and neuropsychiatric disorders, including Parkinson’s disease, schizophrenia, attention-deficit/hyperactivity disorder, autism spectrum disorder and substance use disorder (SUD) [1–6]. DA neurotransmission is tightly controlled by the reuptake of synaptically released DA by the plasma membrane DA transporter (DAT). DAT expressed exclusively in dopaminergic neurons. The proper level of DAT in the plasma membrane of dopaminergic axons is essential for efficient removal of extracellular DA and to control the amplitude and duration of DA neurotransmission. DAT is a major target of psychostimulants, such as amphetamines (Amph) and cocaine, which are substrates and a competitive inhibitor of DAT, respectively. However, mechanisms by which these abused compounds affect DAT function, and its subcellular distribution are not fully understood.

The abuse potential and psychomotor stimulant properties of Amph have been linked to its ability to elevate DA concentrations in the striatum. Resulting increased dopaminergic input in the striatum, mainly in ventral striatum including nucleus accumbens compacta (NAc), has been associated with the rewarding properties of Amph [7–11]. Several mechanisms, such as increased DA release, competitive inhibition of DA reuptake, and conversion of DAT to a substrate efflux mode have been implicated in Amph-induced increase in extracellular DA concentration [10, 12, 13]. Acute (one-time) treatment with Amph has also been shown to reduce DAT surface levels by inducing DAT endocytosis in acute midbrain (MB) slices as measured using slice biotinylation [14], cultured dopaminergic neurons [15], striatal synaptosomes [16, 17], and heterologous cells [18–21]. Various mechanisms have been implicated in Amph-induced DAT endocytosis, including activation of protein kinase C (PKC) [22], Akt [23] and RhoA GTPase [14]. In contrast, our studies using fluorescence and electron microscopy [24] and others [25] observed that acute Amph treatment does not stimulate DAT endocytosis in dorsal striatum (dST) and midbrain in the intact brain and acute brain slices *ex vivo*.

Relatively little is however known about DAT endocytosis and trafficking in the intact brain during repetitive Amph exposure. Repetitive abuse of Amph results in a progressive amplification in the behavioral and neurochemical response, a phenomenon termed sensitization [26]. Sensitization may persist months after administration, thus mimicking long-term sensitivity to drugs as observed in human addicts. At the neurochemical level, sensitization to Amph is manifested in lasting hyper-responsiveness of dopaminergic pathways, which correlates with an increase in the concentration of extracellular DA after a given dose of Amph in rodents [7, 8, 27–30] and humans [31, 32]. In the present study, we explored effects of Amph challenge on DAT in Amph-sensitized DA neurons. To this end, we developed an Amph-sensitization protocol in knock-in mice expressing HA-epitope tagged DAT (HA-DAT) [33]. Analysis using multiple complimentary approaches demonstrated rapid Amph-induced down-regulation of the HA-DAT protein in the striatum of Amph-sensitized mice. This down-regulation was shown to involve endocytic trafficking and was dependent on RhoA effector, Rho-associated kinases (ROCK1/2) activity. Surprisingly, this DAT down-regulation was strictly sex-dependent as it was observed in male but not female animals.

## MATERIALS AND METHODS

### Animals

Male and female (7-8 weeks of age) HA-DAT knock-in mice (on the C57BL/6J background) [33, 34] were used in this study. During experiments, all mice were single housed in cages in regulated environment (23 ± 1°C, 50 ± 5% humidity) on a 12-h light/12-h dark cycle and were fed ad libitum. All procedures were conducted in accordance with the National Institutes of Health’s *Guide for the care and use of laboratory animals* and with the approval of the Institutional Animal Care and Use Committee of the University of Pittsburgh.

### Antibodies and Chemicals

Antibodies were purchased from the following sources: mouse monoclonal antibody against the HA11 epitope (16B12) were from BioLegend (mms-101p), rat monoclonal antibody against the N-terminus of DAT (MAB369) and rabbit polyclonal antibody against tyrosine hydroxlase (TH) (AB152) were from EMD Millipore, Rabbit polyclonal antibodies against vacuolar protein sorting-associated protein 35 (VPS35) were from NovusBio (NB100-1397). Rabbit polyclonal antibodies against vacuolar protein sorting-associated protein 26 (VPS26) was kindly provided by Dr. J. Bonifacino (National Institute of Child Health and Disease). Mouse monoclonal antibody to vesicular monoamine transporter 2 (VMAT2) were from Santa Cruz Biotechnology (sc-374079). Secondary IRDye-800 and IRDye-680-conjugated goat anti-mouse IgG1, anti-rat, and anti-rabbit antibodies were purchased from LI-COR Biosciences; AlexaFluor-488 (A488)- and Cy5-conjugated donkey anti-mouse and anti-rabbit secondary antibodies were from Jackson ImmunoResearch Laboratories. Paraformaldehyde was from Electron Microscopy Sciences. All other reagents and supplies were from Thermo Fisher Scientific or Sigma Chem unless noted otherwise.

### Amph sensitization protocol

*d*-Amphetamine hemisulfate (Amph) (Sigma) was dissolved in normal saline and administered intraperitoneally (i. p.). Amph (0.1 mg/ml) was administered at a dose of 1 mg/kg for amphetamine-induced sensitization by daily injections for 7 consecutive days (Figure 1A). Equivalent volumes of saline were injected in control mice. After the 7-day withdrawal, mice were challenged on day 14 with saline or Amph (1 mg/kg). For behavior experiments, locomotor activity was recorded on days 1, 4, 7 and 14. For biochemical experiments, the mice were euthanized with isoflurane inhalation followed by decapitation, and brains removed on day 7, after withdrawal (7DW) on day 14 before and after 1-hr challenge injections with saline or Amph. Different animals were used for behavior and biochemical experiments. To separate the effects of novelty from the pharmacological effects of the drug, for behavior experiments, animals were acclimated for 3 days (Habituation) to the testing room and locomotor chambers and injected with saline prior to the sensitization protocol. For biochemical experiments, the mice were not habituated to behavior testing room or locomotor chambers but received a saline injection for 3 days of habituation period (Figure 1A).

**Figure. 1.**
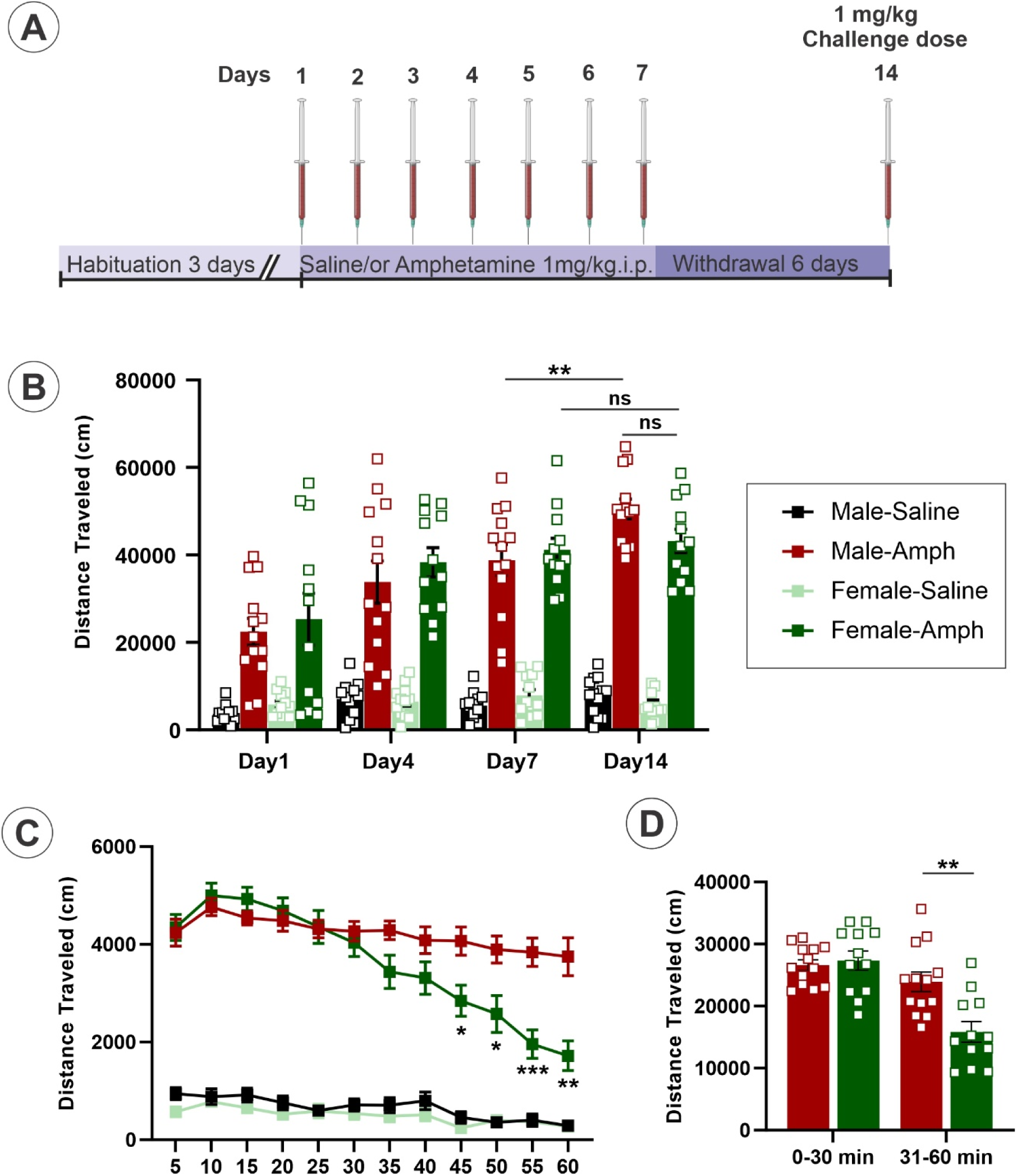
Repeated administration of Amph results in behavioral sensitization in male mice. (A) Schematics of the Amph sensitization protocol. On all days, animals were injected i.p. after 60-min habituation in locomotion chambers. (B) Locomotion activity was measured for 1 hour after saline or Amph injection on day 1,4,7 and after Amph challenge on day 14. Two-way repeated measures ANOVA analysis showed a significant main effect of group (*F*_3,46_ = 70.44, *p*˂0.0001), day (*F*_2.232, 102.7_= 27.90, *p*˂0.0001), and interaction of group x day (*F*_9,138_= 7.729, *p*=0.000). *Tukey’s post hoc* analysis revealed highest locomotor activity on day 7 and 14 of AMPH treated male and female mice compared to respective saline groups (Male: Day 7; p=0.003 and Day 14; p˂0.0001, and Female: p˂0.0001 and p˂0.0001, respectively). (C) Time-course of the locomotion activity during a 1-hr Amph challenge on Day 14. Two-way repeated measures ANOVA analysis showed a significant main effect of group (*F*_3,46_ = 150.4, *p*˂0.0001), day (*F*_3.850, 177.1_= 35.94, *p*˂0.0001), and interaction of group x day (*F*_33,506_= 10.13, *p*=0.000). *Tukey’s post hoc* analysis revealed decrease in the locomotor activity in AMPH challenged female mice compared to male at 45, 50, 55, 60 min. (D) Comparison of the total distance travelled by Amph-sensitized male and female mice for the first 30 minutes and the second 30 minutes of the 1-hr locomotion test. Error bars are SEMs. \**p* < 0.05, \*\**p* < 0.01 \*\*\**p* < 0.001, \*\*\*\**p* < 0.0001.

### Locomotor activity

The behavioral tests were conducted in the Rodent Behavior Analysis Core of the University of Pittsburgh Schools of Health Sciences. Locomotor activity (distance travelled) was quantified using an open field arena (40 × 40 × 40 cm; Omnitech Electronics) in a sound-attenuated and environmentally-controlled chamber (60 x 64 x 60 cm; Omnitech Electronics) equipped with 4 horizontal 16 x 16 arrays of infrared photobeam sensors. Measurements were conducted in 5-min bins for 60 min after Amph or saline injection on experimental days 1, 4, 7, and 14, by using behavioral tracking software (Fusion, Omnitech Electronics). On each test day, animals were acclimated to individual activity chambers for 60 min to allow the animal to become accustomed to its behavioral cage before subsequent injections of either Amph (1 mg/kg, i.p.) or saline. Following each injection, the mice were placed back into their respective activity chamber and their locomotor activity was recorded.

### Striatal synaptosome preparation

The synaptosomes were prepared as described [35]. Briefly, striatal tissue was rinsed in ice-cold Gey’s balanced salt solution with 10 mM D-glucose. Striatum was homogenized in 1.5 ml ice-cold HEPES buffer (5 mM HEPES, 0.32 M sucrose, pH 7.4) using a glass homogenizer. The homogenate was centrifuged at 1000xg for 10 min at 4°C, and the resulting supernatant was centrifuged at 12,500xg for 20 min at 4°C to pellet the synaptosomes. Pellets were resuspended in Krebs-Ringer HEPES buffer (KRH; 120 mM NaCl, 4.7 mM KCl, 2.2 mM CaCl_2_, 1.2 mM MgSO_4_, 1.2 mM KH_2_PO_4_, 10 mM glucose, 10 mM HEPES, pH 7.4) and used for Western blot analysis or DA uptake parameter measurements.

### Synaptosome DA Uptake

Synaptosomes from two mice for each individual experiment and pellets were resuspended in KRH. Synaptosomes were incubated with 20 nM [^3^H]DA (Perkin Elmer Life Sciences, Boston, MA) and unlabeled DA (0.031, 0.0625, 0.125, 0.25, 0.5 and 1.0 µM) for 10 min at 37 °C in KRH supplemented with 10 µM ascorbic acid, 10 µM pargyline, 1 µM desipramine and 10 µM cathechol-O-methyl-transferase inhibitor. Nonspecific [^3^H]DA accumulation was determined in the presence of 100 µM cocaine. The reaction was terminated by adding ice-cold KRH and immediate washing with KRH by filtration through Whatman GF/A glass fiber filters using a Brandel cell harvester (Model M-48 Biochemical Research and Development Laboratories Inc., Gaithersburg, MD). Filters were then solubilized in 0.1N NaOH/1% SDS. Synaptosomes-associated [^3^H]DA was measured by liquid scintillation counting. Kinetic parameters (K_m_ and V_max_) for [^3^H]DA uptake were calculated using Michaelis-Menten equation by GraphPad software.

### Western blotting protein detection

The mice were euthanized, and the brains were rapidly extracted and sliced into coronal sections (0.4 mm thick) using a brain matrix followed by either preparation of synaptosomes from striatum as described above or extraction of tissue from specific brain regions. Tissues from dStr, NAc, Ventral tegmental area (VTA), and Substantia nigra compacta (SNc) were extracted using a 1.0-mm Harris Uni-Core micropunch (Electron Microscopy Sciences, Mountain Lakes Blvd, Redding, CA, USA). Synaptosomes and brain tissues were solubilized in the lysis buffer containing 50 mM Tris-HCl pH 7.4, 0.5 % sodium deoxycholate, 1% NP-40, 150 mM NaCl, 2mM EGTA, 2mM EDTA and phosphatase and protease inhibitors. Lysates were cleared by centrifugation for 15 min at 16,000 x g at 4^0^C. Aliquots of lysates (30 µg protein for Striatal synaptosomes, dST, and NAc and 50 µg for whole MB, VTA and SNc, respectively) were denatured in sample buffer at 37^0^C for 30 min. Aliquots of brain tissue lysates were resolved by 7.5 % SDS-PAGE, transferred to nitrocellulose (Li-COR) and probed with appropriate primary and secondary antibodies conjugated to far-red fluorescent dyes (IRDye-680 or -800) followed by detection using Odyssey Li-COR system. Quantifications were performed using ImageJ software.

### Immunofluorescence staining

Mice were injected intraperitoneally with xylazine/ketamine, and perfused transcardially with 50 ml of PBS and 50 ml of 4% paraformaldehyde (PFA) in PBS (pH 7.2). Brains were then post-fixed with 4% PFA for 4 hrs at 4 °C and cryoprotected in 20% sucrose/TBS overnight followed by 30% sucrose in PBS at 4°C until ready for cryosectioning. Brains were embedded in OCT compound (Sakura Finetek), and deep frozen in liquid nitrogen. Sagittal free-floating brain sections were made at 50 µm thickness using freezing microtome (Leica CM 1950). Sections were incubated in PBS containing 1% H_2_O_2_ for 15 min, permeabilized with 0.1% Triton X-100 for 1 h, and preincubated with blocking buffer containing 10% normal donkey serum (D9663, Sigma Millipore), 3% BSA (A2153, Sigma Millipore), 0.1% Triton X-100 (Sigma Millipore) in PBS for 1 h at room temperature. After blocking, sections were incubated with mouse HA11 (1:500) and rabbit VPS35 (1:500) at 4°C in blocking solution for 48 hrs and washed, followed by donkey anti-mouse conjugated to Alexa488 and Cy5-conjugated donkey anti-rabbit secondary antibodies for 1 hr at room temperature. Nuclei were stained with Hoechst 33342 (62249, Thermo Fisher Scientific). Sections were mounted in Prolong Gold antifade mounting medium (P36930, Thermo Fisher Scientific).

### Confocal microscopy and quantification of co-localizations

High-resolution z-stack of confocal images of DA neurons in sagittal sections were acquired using a spinning disk confocal Marianas system (Intelligent Imaging Innovation, Denver, CO) as described [36]. Typically, 20-30 serial two-dimensional confocal images of cryosections were recorded at 400 nm intervals. All image acquisition settings were identical for all experimental variants in each experiment.

To quantify the amount of HA-DAT colocalized with VPS35, 3D images were cropped to generate new images consisting of a *z* stack of three consecutive confocal sections away from the section corresponding to the surface of a slide displaying strong labeling and chosen by a similar extent of fluorescence intensity (antibody penetration). These images were deconvolved using a No Neighbors algorithm of SlideBook 6. An automated segment mask was generated from background-subtracted images to select voxels detected through the 488 nm channel (Mask 1; HA-DAT) and the 640 channels (Mask 2; VPS35). A “colocalization” mask (Mask 3) was generated to select voxels positive for both 488 nm and 640 nm channels. The sum fluorescence intensity of the 488 nm channel in the colocalization mask was divided by the sum fluorescence intensity of HA-DAT (Mask 1) to calculate the fraction of HA-DAT colocalized with VPS35.

### Statistical Analysis

All statistical analyses were performed using GraphPad Prism software (GraphPad). For comparisons of each two groups, unpaired Student’s *t* test and for comparisons of more than two groups, one way ANOVA followed by Tukey’s test was used. For behavior study, a two-way repeated measure ANOVA followed by Bonferroni ’s multiple comparison tests was used. All experiments were performed at least three times. Differences were considered significant when the *p* value was <0.05, with the specific *p* values detailed within each figure legend.

## RESULTS

### Amph-induced locomotor sensitization

To test whether repeated administration of Amph results in sensitization, we developed a schedule of Amph administration which includes repeated injections and withdrawal. HA-DAT mice were injected with saline or Amph (1 mg/kg, i.p) for 7 days followed by 7 days of withdrawal and then challenged with saline or a single Amph dose (1.0 mg/kg, i.p.) on Day 14 (Figure 1A). Because female and male rodents were previously shown to respond differently to Amph [37–39], we performed these experiments in mice of both sexes. Development of the behavior sensitization was evident by the highest locomotor activity measured during 1-hr Amph challenge on Day 14 in both sexes (Figure 1B-C; two-way ANOVA for repeated measures indicated significant effect of group (*F*_3.46_= 70.44, *p*˂0.0001), day (*F*_2.232, 102.7_= 27.90, *p*˂0.0001), and interaction of group x day (*F*_9,138_ = 7.729, *p*=0.0001). Tukey’s test revealed that the Amph-treated male and female mice were significantly different from the saline-treated mice. However, the response was significantly stronger on day 14 compared to day 7 in Amph-treated male but not female mice (Figure 1B). Interestingly, the time-course of the distance traveled during Day 14 challenge demonstrated a high initial activity in both sexes followed by the statistically significant reduction of this activity after 30 min in female but not male mice (Figure 1C-D). Such sex-specific behavior kinetics was not observed after single doses of Amph at earlier time points of the sensitization schedule, although the behavior stimulation by acute Amph was evident (Figure S1).

### Amph challenge in Amph-sensitized mice triggers DAT protein down-regulation in the striatum in sex-dependent manner

To investigate mechanisms underlying behavior sensitization described in Figure 1, we examined whether Amph sensitization affects DAT, a principal transporter that Amph uses to enter dopaminergic neurons. Hence, DAT protein levels were measured by western blotting with the antibody recognizing the amino-terminal region of DAT in striatal synaptosomal preparations and MB homogenates obtained on Day 14 from saline- or Amph-challenged male and female mice. The level of tyrosine hydroxylase (TH) did not change during Amph sensitization and challenge, indicating that Amph administration did not have a general toxicity effect on dopaminergic neurons. Therefore, TH immunoreactivity was used as a normalization factor for DAT immunoreactivity. HA-DAT protein level in striatal synaptosomes in Amph-sensitized male mice was found to be significantly reduced compared to saline-treated mice (t-test: t = 3.9414, df = 20, p = 0.0008) (Figure 2A-B). By contrast, no difference in the HA-DAT amount was detected between saline and Amph-challenged female mice (t-test: t = 0.8994, df = 18, p = 0.3803) (Figure 2A-B). To rule out the possibility of Amph-induced conformational changes or post-translational modifications interfering with the recognition of the amino-terminal domain of DAT by the antibody, HA-DAT down-regulation in male was confirmed using HA antibody (Figure S2A). The amount of HA-DAT in MB lysates was not affected by Amph challenge in sensitized male (t-test: t = 0.4704, df = 20, p = 0.6431) and female mice (t-test: t = 0.3718, df = 18, p = 0.7144) (Figure 2D-E). The amount of vesicular monoamine transporter 2 (VMAT2), a transporter that is responsible for accumulation of Amph in synaptic vesicles of dopaminergic neurons, was not affected in striatal synaptosomes (Figure 2A and C) and MB lysates by Amph challenge (Figure 2D and F). These data demonstrate a high level of DAT specificity of Amph effects in sensitized male mice.

**Figure. 2.**
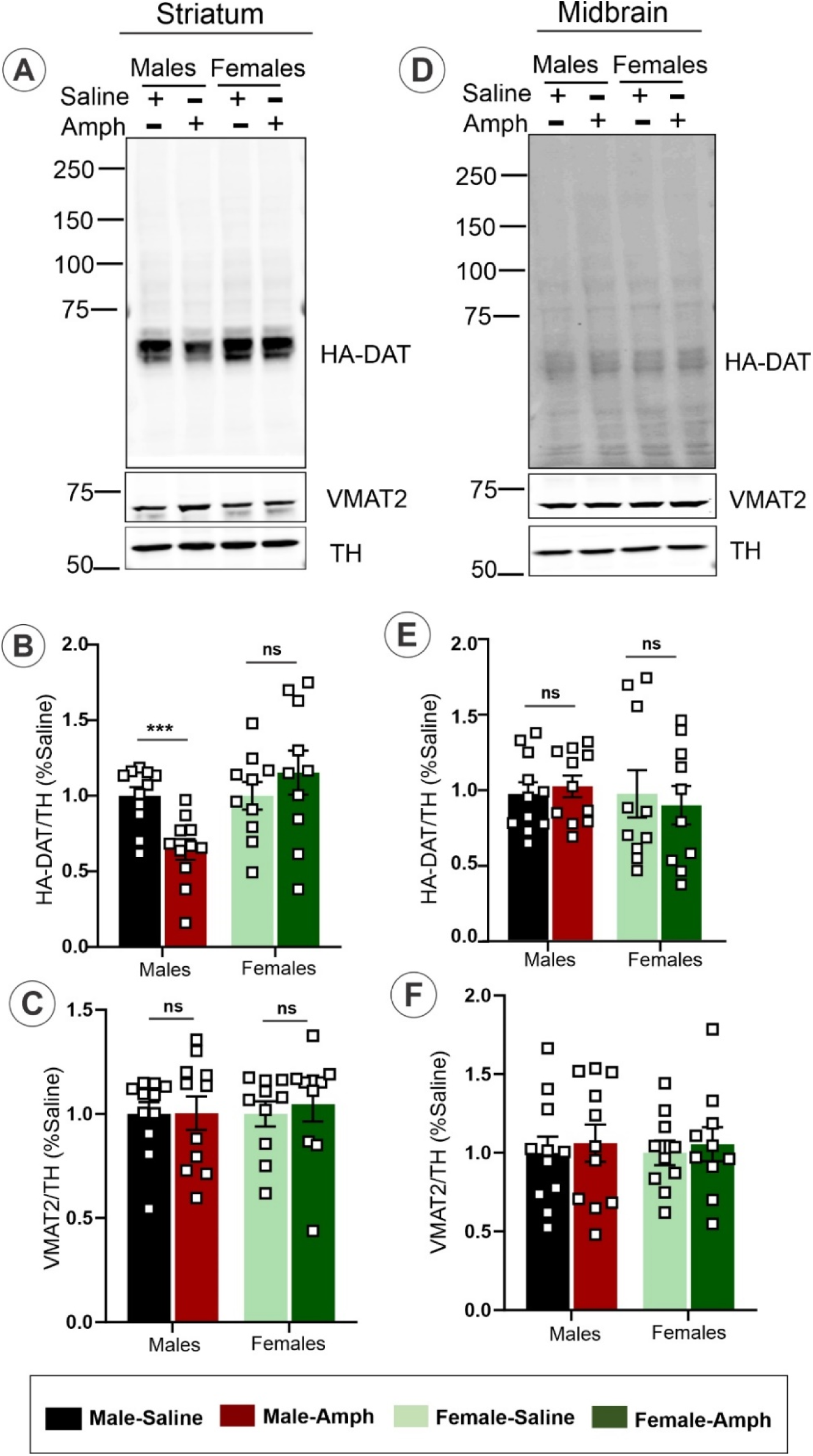
Amph challenge in Amph-sensitized mice decreases the DAT protein level in striatum in sex-dependent manner. HA-DAT mice sensitized according to the protocol in Figure 1A were challenged with saline and Amph for 1 hr on Day 14. Striatal synaptosomes [30 µg/lane] (**A**-**C**) and midbrain tissue [50 µg/lane] (**D-F**) were lysed, and the aliquots of lysates **(C, F)**] were resolved by 7.5 % SDS-PAGE, respectively, transferred to nitrocellulose and probed with rat DAT, rabbit TH and rabbit VMAT2 antibodies by Western blotting. Representative blots are shown. Bar graphs show the mean values (± S.E.M) of the amounts of HA-DAT (**B** and **E**) or VMAT2 (**C** and **F**) expressed per brain as fractions of the amounts in saline samples and normalized by the amount of TH from 3 and more independent experiments. Asterisks indicate significant differences of “Amph” samples compared to Saline groups, p ˂ 0.05 (Student’s unpaired t-test [n = 5]). *ns,* no significant difference.

To confirm that HA-DAT down-regulation in striatal synaptosomes on day 14 was caused by the Amph challenge in Amph-sensitized male mice, we measured HA-DAT expression levels in Amph-injected mice on Day 7 and Day 14 after withdrawal but before the 1-hr Amph challenge (Figure S2B). Western blot analysis showed no difference in the amount of HA-DAT on Day 7 (t-test: t = 0.8890, df = 10, p = 0.3949) and after withdrawal (t-test: t = 0.5611, df = 10, p = 0.5871) in Amph-sensitized male mice compared to saline-treated mice (t-test: t = 4.5808, df = 10, p = 0.001). In the same group of male mice, DAT protein level was significantly reduced following the Amph challenge (t-test: t = 4.5808, df = 10, p = 0.001) as compared to that level in Amph-sensitized after withdrawal and in saline-treated sensitized mice. Altogether, the data in Figure 2 and Figure S2 demonstrate that Amph-challenge results in down-regulation of the axonal DAT protein in sensitized male and not female mice.

### Amph-induced DAT down-regulation involves endocytic trafficking

We and others have previously shown that the bulk of DAT is located at the surface of dopaminergic axons in striatum whereas a pool of intracellular DAT in these axons is extremely small [24]. Therefore, Amph-induced down-regulation of HA-DAT in striatum described in Figure 2 must be associated with the re-distribution of a substantial fraction of DAT from the cell surface to endosomes and/or lysosomes via endocytosis. To determine whether Amph challenge of sensitized mice leads to down-regulation of the cell-surface HA-DAT (endocytosis) in sex-dependent manner, [^3^H]DA uptake assays in synaptosomal preparations were performed and kinetics parameters were measured (Figures 3A-C). Amph challenge reduced the maximal DA uptake velocity (Vmax) by ∼35-45% in Amph-treated males (Amph: Vmax = 50.9 ± 8.4 versus control: Vmax = 80.9 ± 9.8; P = 0.0157), whereas no change in the same parameter was observed in experiments with Amph-challenged female mice (Amph: Vmax = 40.8 ± 2.2 VS control: Vmax = 43.5 ± 3.6; P = 0.3286) (Figures 3A-C). By contrast, the K_m_ values of the DA uptake were not significantly different among all experimental variants, suggesting that sensitization and Amph challenge do not affect the substrate affinity of DAT. Thus, the observation of decreased Vmax is consistent with the hypothesis whereby Amph challenge induces DAT endocytosis leading to the decrease in DAT concentration at the surface of dopaminergic axons.

**Figure. 3.**
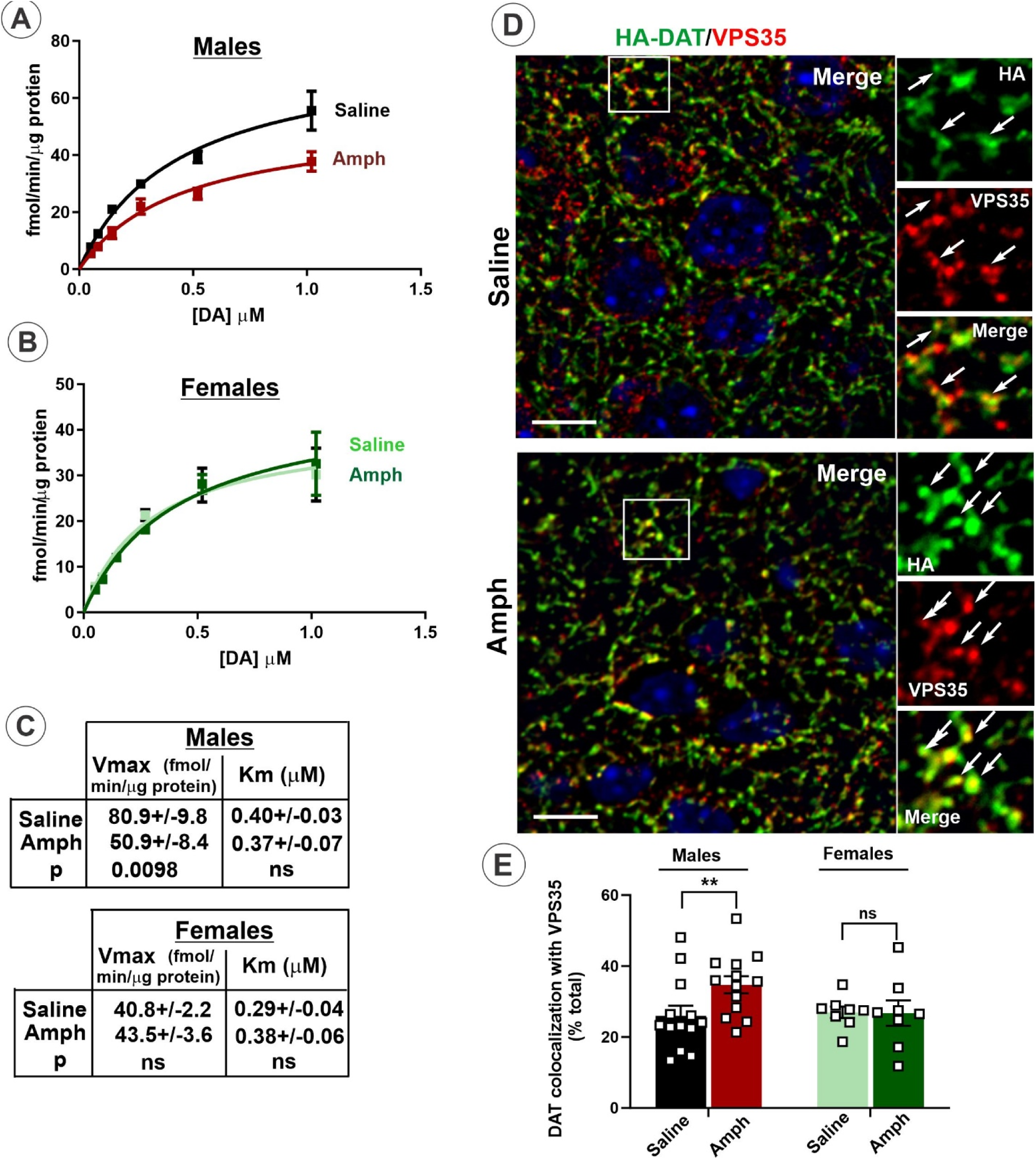
Amph challenge of sensitized male mice exhibit reduced density of HA-DAT at the cell surface and increased DAT co-localization with VPS35 in striatal axons. Kinetic analysis of [^3^H]DA uptake was performed in striatal synaptosomes of male (**A**) and female (**B**) mice that were challenged with saline or Amph on day 14. [^3^H]DA (20 nM) and increasing concentrations of unlabeled DA (from 0.005 to 1 µM) were applied simultaneously. Mean values (± SEM) of kinetic parameters (maximal velocity Vmax and affinity Km) derived from experiments exemplified in **A** and **B** are presented in (**C**). Data are from three separate experiments, each performed in triplicates. (D) Brains were isolated 1 hour after saline of AMPH challenge on Day 14 and fixed by cardiac perfusion with 4% paraformaldehyde. Sagittal cryosections were permeabilized and labeled with antibodies against HA (HA-DAT, *green*) and VPS35 (*red*) followed by fluorophore-conjugated secondary antibodies. Nuclei were stained with DAPI (*blue*). 3-D images of the striatum were acquired by spinning disk confocal microscope. Maximum-intensity projections of three consecutive confocal sections from the collected z-stacks are shown. Insets show high-magnifications images of areas indicated by white rectangles. Arrows point to examples of HA-DAT/VPS35 co-localization in axonal varicosities. Scale bars, 10 µm (E) Quantification of co-localization of HA-DAT and VPS35. Bar graphs show mean values (± S.E.M) of the percent of HA-DAT co-localized with VPS35 of the total HA-DAT fluorescence in 3-D images exemplified in A. **p ˂ 0.01; ns, no significant difference (Student’s unpaired t-test).

To further investigate the Amph-induced endocytic trafficking process leading to DAT down-regulation, we analyzed HA-DAT localization in striatal axons. We and others recently demonstrated that a pool of axonal DAT is located in endosomes containing VPS35, a subunit of the retromer complex mediating recycling from endosomes, whereas markers of sorting and late endosomal compartments were undetectable in dopaminergic axons [24, 36, 40]. Therefore, co-labeling of HA-DAT and VPS35 was performed on sagittal cryosections of the mouse brain. A substantial colocalization of HA-DAT with VPS35 was observed mainly in axonal varicosities (synaptic areas) in the striatum of both sexes (Figure 3D). Quantifications revealed that Amph challenge resulted in a significant increase of HA-DAT co-localization with VPS35 in striatal axons in males (t-test: t = 2.3475, df = 24, p = 0.0275) but not in females (t-test: t = 0.0598, df = 14, p = 0.9532) (Figure 3E). The amount of VPS35 and another retromer complex subunit VPS26 in striatum and midbrain did not significantly change among all experimental variants (Figure S3). Together, data in Figure 3D-E further support the notion that Amph challenge redistributes a fraction of HA-DAT from the plasma membrane to endosomes in a subset of DA axons in the striatum. Certainly, diffraction-limited resolution of confocal microscopy does not allow formal determination of whether HA-DAT immunofluorescence co-localized with VPS35 informs about endosomal HA-DAT given a small size of dopaminergic presynaptic areas. Furthermore, background fluorescence of brain tissue imaging makes the use of the 3D super-resolution imaging and its quantitative analysis to be technically challenging. Nevertheless, together with the observation of Amph-induced decrease in Vmax (Figure 3A-C), increased co-localization of HA-DAT with retromer in males indicates that HA-DAT is internalized and accumulated in early/recycling endosomes after Amph-challenge-induced endocytosis.

To examine whether downregulation of the DAT protein measured in striatal synaptosomes of male mice involves traffic of internalized DAT through the endolysosomal system, Amph-sensitized mice were pretreated with chloroquine (CQ; 10 mg/ml; i. p.). CQ is a membrane-permeable base compound that accumulates in acidic compartments, such as endosomes, autophagosomes and lysosomes, which results in neutralizing pH in these compartments and blocking endosomal maturation and lysosomal degradation processes. CQ did not significantly influence the amount of HA-DAT in saline-treated mice, but decreased Amph-induced DAT down-regulation in striatal synaptosomes by ∼46% (Amph+CQ group compared to the control “Amph” group) (Figure 4A). These data demonstrate the involvement of DAT traffic through endosomal and lysosomal compartments in Amph-induced DAT down-regulation.

**Figure 4.**
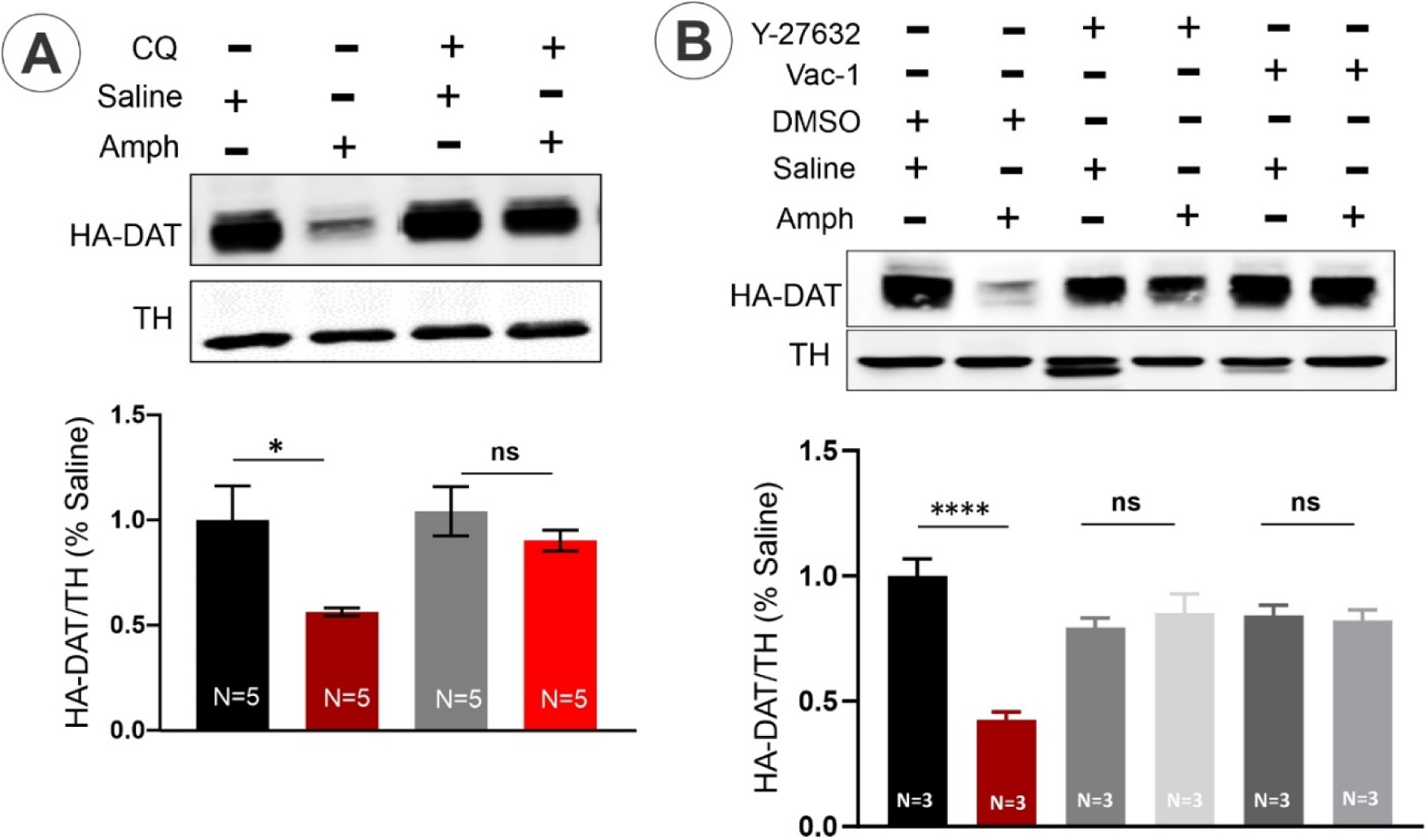
Amph-induced down-regulation of HA-DAT in sensitized mice in striatum is blunted by inhibitors of endolysosomal traffic and ROCK1/2. (A) Chloroquine (CQ) (100 mg/kg) (“+”) or vehicle (“-“, saline) were injected i. p. 2 hours before saline or Amph challenge of male mice on Day 14 of the sensitization protocol. Striatal synaptosomes were prepared 1 hour after saline/Amph challenge injections, electrophoresed and probed by immunoblotting with antibodies to DAT and TH (loading control). Representative immunoblot is shown. Bar graphs show mean values (+/-SEM) of the amounts of HA-DAT normalized to the amount of TH and expressed as fractions of the TH-normalized amounts of HA-DAT in “Saline” samples. (B) Vehicle (DMSO, “-“), ROCK1/2 inhibitor Y-27632 (5 mg/kg) or vacuolin-1 (2 mg/kg) were injected i. p. 2 hours before saline or Amph challenge of male mice on day 14 of the sensitization protocol. Striatal synaptosomes were prepared 1 hour after saline/Amph challenge injections, electrophoresed and probed by immunoblotting with antibodies to DAT and TH (loading control). Representative immunoblot is shown. Bar graphs show mean values (+/-SEM) of the amounts of HA-DAT normalized to the amount of TH and expressed as fractions of the TH-normalized amounts of HA-DAT in “Saline” samples. One-way ANOVA, P < 0.0001, Tukey’s multiple comparisons test, **** P < 0.0001, *** P < 0.001 and * P < 0.05. Error bars represent SEMs. Numbers of animals are indicated on bars.

Further inhibitory analysis was performed to better define mechanisms of endocytic DAT down-regulation. Studies by Wheeler and co-workers demonstrated that acute Amph induces DAT endocytosis involving activation of RhoA GTPase [14]. In this study, DAT endocytosis was shown to be sensitive to inhibitors of the main downstream effectors of active RhoA, the Rho-associated coiled-coil containing kinases (ROCK1/2) [41]. Intraperitoneal injection of the ROCK1/2 inhibitor Y-27632 was previously shown to result in ROCK1/2 activity inhibition in mouse brain [42]. Therefore, we tested the effect of Y-27632 on HA-DAT down-regulation induced by Amph challenge in male mice and found that this inhibitor dramatically attenuates this Amph effect (Figure 4B). This result suggested that DAT internalization via a RhoA-mediated pathway is involved in observed DAT down-regulation.

Autophagic degradation of DAT was implicated in the effects of the injection of cocaine in dorsal striatum (dStr) on DAT, although endocytosis and localization of DAT in early phagophores or autophagosomes after cocaine injection has not been demonstrated [43]. The latter study used vacuolin-1 as an inhibitor of autophagy to support their model. However, vacuolin-1 has been shown to impose wide-range effects on endosome-lysosome system through the inhibition of PIKfyve kinase located in early, sorting and late endosomes, and also through increased activity of Rab5 that blunts endosome maturation [44–46]. In our experiments, vacuolin-1 also blocked Amph-induced down-regulation of HA-DAT (Figure 4B), further supporting the role of post-endocytic traffic of DAT in Amph-challenged sensitized male mice. While the possibility of autophagocytic processes in dopaminergic synapses has been proposed [43, 47, 48], the direct evidence (visualization of phagophores or autophagosomes) supporting autophagy in presynaptic areas in intact brain has not been provided. Studies in intact brain did not detect lysosomal compartments in dopaminergic axons [24, 36, 49, 50]. Further, the essential macroautophagy protein, ATG9, was not detected by immunofluorescence staining of dopaminergic axons in HA-DAT mice, although it was readily detected in non-dopaminergic cells in the striatum and in the soma of dopaminergic neurons (Figure S4). Therefore, we suggest that upon Amph challenge, DAT is trafficked and possibly degraded through conventional endocytic trafficking mechanisms rather than via autophagy.

### Amph-induced DAT downregulation in sensitized male mice is brain-region-specific

Our fundings of DAT endocytic trafficking triggered by the Amph challenge of sensitized males (Figures 2-4) are somewhat inconsistent with our previous morphological studies which demonstrated minimal constitutive and acute Amph-induced endocytosis of DAT in dStr [24]. Interestingly, DAT down-regulation triggered by activation of protein kinase C (PKC) was observed using biotinylation of acute slices in NAc but not in dStr, suggesting higher DAT endocytic traffic activity in NAc than in dStr [51, 52]. Region-specific endosomal localization of DAT within striatum is technically difficult to analyze using light microscopy of sagittal slices. Therefore, to test whether Amph effects on HA-DAT in sensitized mice are brain-region specific, immunoblotting analysis of HA-DAT in tissue lysates of NAc, dST, VTA, and SNc isolated from saline- or Amph-challenged sensitized mice was performed. This analysis revealed that Amph-induced DAT down-regulation occurs in NAc (t-test: t = 2.8219, df = 10, p = 0.0181) (Figure 5A) rather than in dStr (Figure 5B). No changes in HA-DAT levels were observed in VTA (Figure 5C), and SNc (Figure 5D) recovered from male mice. No significant difference in the amount of HA-DAT in Amph-sensitized compared to control was observed in either of the brain areas of female mice (Figure 5).

**Figure 5.**
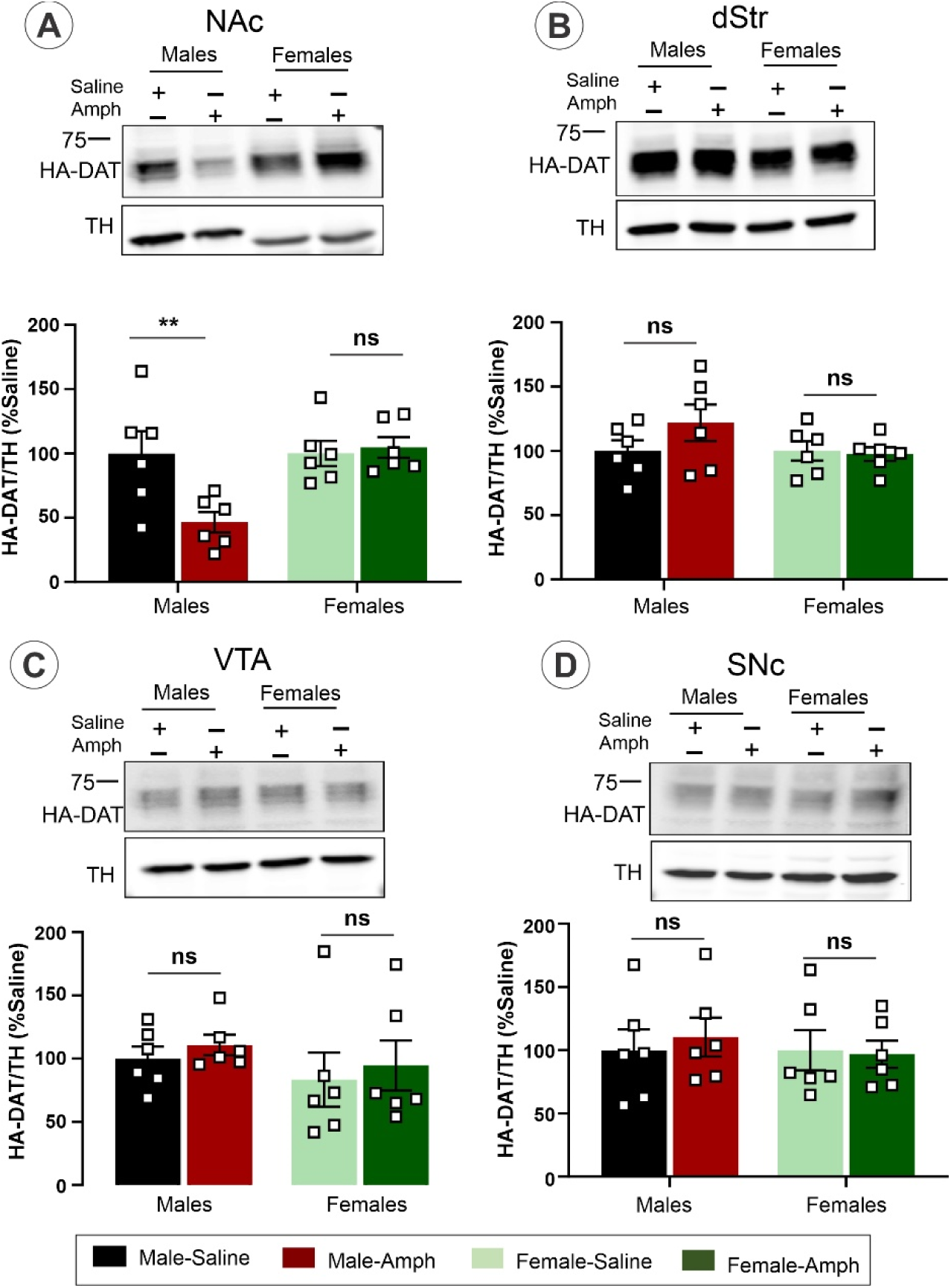
Brain-region specific downregulation of HA-DAT in Amph-sensitized male mice. Tissues from NAc (**A**), dStr (**B**), VTA (**C**), and SNc (**D**) were isolated from sensitized mice challenged with saline or Amph on Day 14. Lysates were electrophoresed and probed by western blotting with antibodies to DAT and TH. Representative DAT and TH immunoblots are shown. Graph bars are mean values of the amounts of HA-DAT normalized to the amount of TH and expressed as percent of the mean TH-normalized amounts of HA-DAT in “Saline” samples in each individual experiment. Analysis revealed that HA-DAT protein down-regulation occurs significantly in NAc (t-test: t = 2.8219, df = 10, p = 0.0181) (**A**) but not in dStr (t-test: t = 1.3371, df = 10, p = 0.2108) (**B**), VTA (t-test: t = 0.8602, df = 10, p = 0.4098) (**C**), and SNc (t-test: t = 0.4611, df = 10, p = 0.6546) (**D**) in Amph-sensitized male compared to control (“saline”) mice. No significant differences were observed in NAc (t-test: t = 0.3694, df = 10, p = 0.7195) (**A**), dStr (t-test: t = 0.2559, df = 10, p = 0.8032) (**B**), VTA (t-test: t = 0.1256, df = 10, p = 0.9025) (**C**), and SNc (t-test: t = 0.1640, df = 10, p = 0.8730) (**D**) in Amph-sensitized female compared to control mice.

## DISCUSSION

In this study, we have used several complimentary approaches to demonstrate that a short Amph challenge of Amph-sensitized mice results in a substantial reduction in the DAT protein in the striatum via DAT endocytic trafficking in males but not female mice. The observations of sex-dependent DAT down-regulation (Figures 2, 5 and S2A) are consistent with the differences in the time course of Amph-induced locomotor activity in males and females (Figure 1E). In females, DAT is not down-regulated, and therefore, an excess of DA is removed, which results in the temporary increase followed by the decrease in the locomotor activity; in males DAT is down-regulated, extracellular DA is not efficiently removed, and a high locomotor activity is sustained.

Most previous studies in rodents show a more robust behavioral sensitization of females than males in response to single or repeated injections of amphetamines [53], although there have been reports of a greater Amph-induced sensitization in males [54]. However, direct comparison of these studies with our results demonstrating that female mice do not exhibit a sustained Amph-induced locomotor sensitization is difficult. First, much higher doses of Amph compared to our experiments have been used in previous studies (for example, [53]). Second, most studies performed 30-min tests [55–57] which may have precluded detection of the different behavior kinetics in males and females observed in our experiments after 30 min of the Amph challenge.

In summary, the data prompt us to hypothesize that Amph challenge of sensitized male mice results in DAT endocytosis in both dStr and NAc, whereas apparent degradation of the DAT protein occurs mainly in NAc. Whether a fraction of DAT is indeed degraded within 1 hr after Amph administration or transported out of striatum, specifically out of NAc, remains unclear. Against the former possibility is the lack of degradative compartments (lysosomes) in the striatal dopaminergic axons as shown by immunofluorescence and electron microscopy studies [24, 36, 49, 50], although morphological approaches are not sensitive to formally rule out the presence of a small number of lysosomes in these axons, especially in a less studied NAc. The latter model would entail rapid microtubular-mediated transport of DAT in endosomes or early autophagocytic compartments to the somatodendritic region of neurons where DAT can be degraded in lysosomes. Our previous analysis of the long-range transport of DAT between striatum and midbrain through medial forebrain bundle (MFB) demonstrated the prevalence of a slower transport mechanism, lateral membrane diffusion [36]. This makes it to be unlikely that a significant number of DAT molecules can be transported through MFB within 1 hour of Amph challenge to allow their lysosomal degradation in the soma. However, we have also detected a subset of MFB axons in which a significant amount of DAT was present in endocytic vesicles [36], suggesting that a fraction of DAT can be transported through MFB using a faster microtubular-dependent mechanism. Future mechanistic analysis of DAT trafficking in NAc would require the development of new methods allowing analysis of DAT trafficking in intact brain to elucidate the molecular mechanisms of Amph-induced DAT down-regulation and its sex-dependence.

## Acknowledgments

We are grateful to Dr. Juan Bonifacino (NICHD) for the generous gift of VPS26 antibody.

## Author Contributions

T.R.B., conceived the experimental strategy; designed experiments; performed experiments; analyzed data; wrote the first draft of the manuscript. A.S., provided funding and experimental model; designed experiments; edited the manuscript.

