## Supplemental figures for "Endocytic down-regulation of the striatal dopamine transporter by amphetamine in sensitized mice in sex-dependent manner"

**Bagalkot and Sorkin**

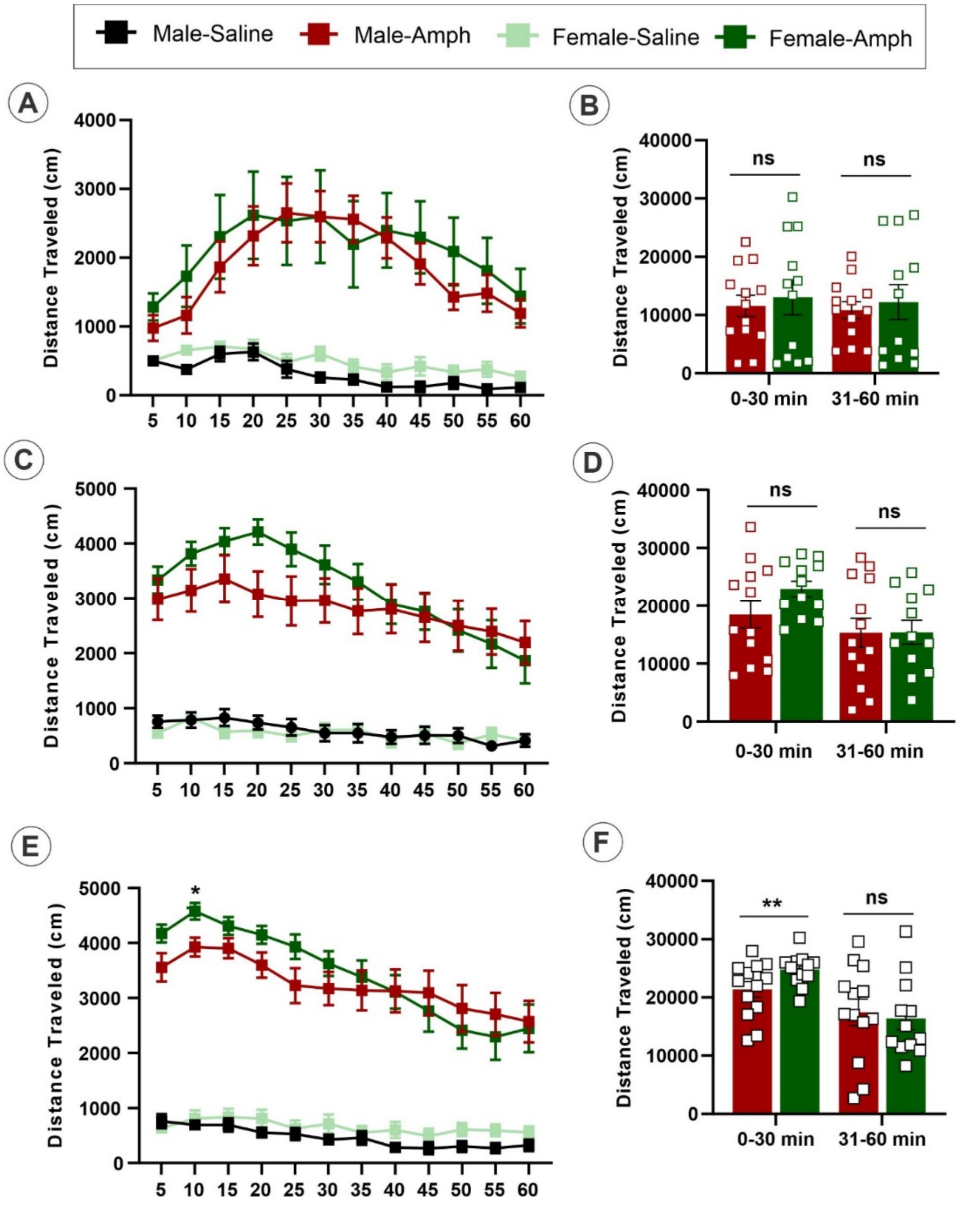


**Figure S1. Time-course of the locomotor activity during a 1-hr Amph injection at different days of the sensitization protocol.**

(**A**) ***Day 1***. Two-way repeated measures ANOVA analysis showed a significant main effect of group ($F_{3,46}$ = 11.42, $p$<0.000), time ($F_{3.333,153.3}$ = 13.69, $p$<0.000), and interaction of group x time ($F_{33,506}$ = 4.503, $p$<0.000). *Tukey's post hoc* analysis revealed increase in locomotor activity in Amph-challenged male and female mice compared to their respective controls.

(**C**) ***Day 4***. Two-way repeated measures ANOVA analysis showed a significant main effect of group ($F_{3,46}$ = 30.29, $p$<0.000), time ($F_{4.742,\ 218.1}$= 27.34, $p$<0.000), and interaction of group x time ($F_{33,506}$ = 6.438, $p$<0.000). *Tukey's post hoc* analysis revealed increase in locomotor activity in Amph-challenged male and female mice compared to their respective controls.

(**E**) ***Day 7***. Two-way repeated measures ANOVA analysis showed a significant main effect of group ($F_{3,46}$ = 66.35, $p$<0.000), time ($F_{3.740,\ 172.1}$= 30.87, $p$<0.000), and interaction of group x time ($F_{33,506}$ = 6.210, $p$<0.000). *Tukey's post hoc* analysis revealed increase in locomotor activity in Amph-challenged male and female mice compared to their respective controls**.**

(**B, D** and **F**) 1 hour locomotion test: the total distance travelled by Amph-sensitized male and female mice for the first 30 minutes and the second 30 minutes at corresponding days of the sensitization protocol**.** Error bars are SEM. *$p$ < 0.05, **$p$ < 0.01 indicates significant difference between amphetamine sensitized male and female mice. *ns* indicates no significant difference.

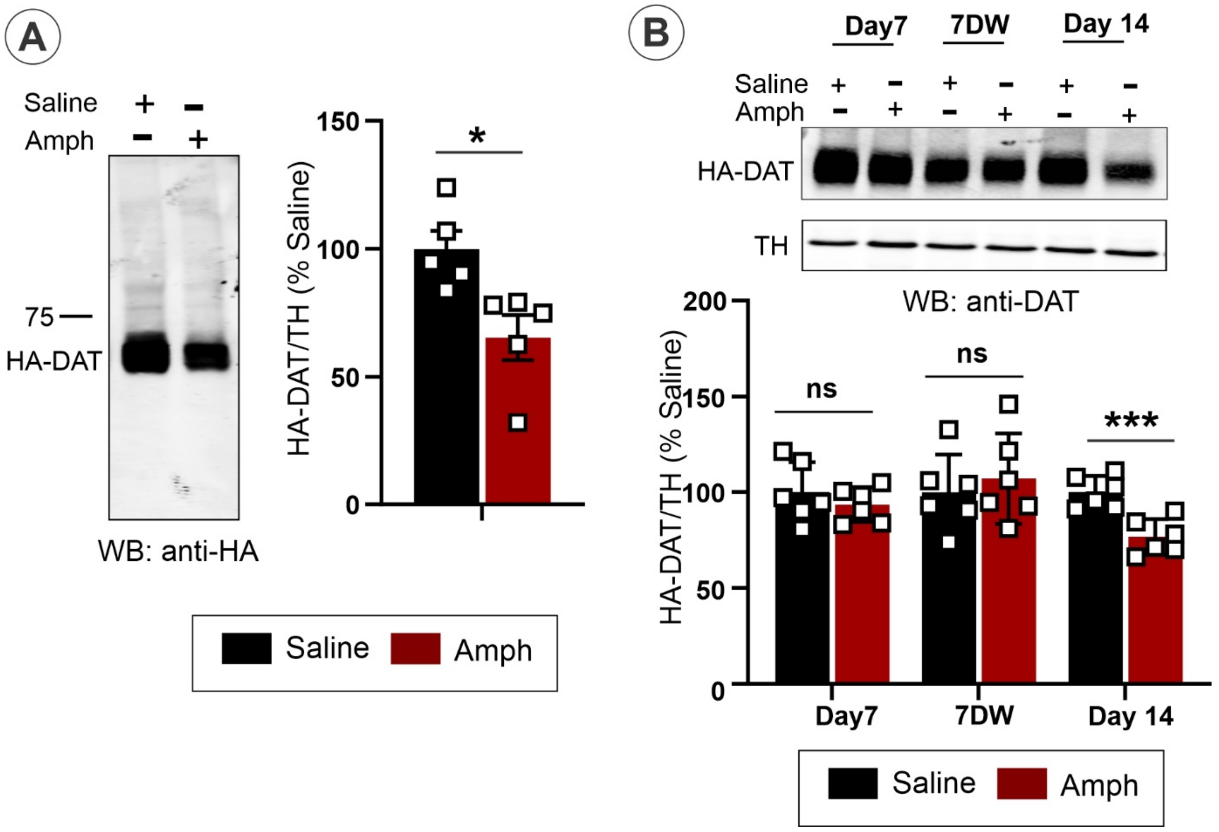


**Figure S2. Amph challenge results in reduced HA-DAT protein levels in striatal synaptosomes only on Day 14 of the sensitization protocol**

(**A**) Striatal synaptosomes obtained after saline and Amph-challenged on Day 14 from male mice were lysed as in Figure 2A. The protein samples [60 µg/ lane for DAT from striatal synaptosomes] were resolved by 7.5% SDS-PAGE, transferred to nitrocellulose and probed with HA and TH antibodies. Representative blot is shown. The bar graph shows the mean (± S.E.M) intensity of bands from 3-4 independent experiments. The amounts of HA-DAT were normalized to the amounts of TH. Asterisks indicate significant differences compared to Saline group, $p < 0.05$ (Student's unpaired t-test [n = 5]).

(**B**) Striatal synaptosomes obtained on Day 7, Day 14 before saline and Amph challenge (*7DW*) and after saline and Amph challenge on Day 14 (*Day 14*) were lysed. The protein samples [30 and 60 µg/ lane for DAT from striatal synaptosomes were resolved by 7.5% SDS-PAGE, transferred to nitrocellulose and probed with DAT and TH antibodies. Representative blot is shown. The bar graph shows the mean ± S.E.M. Intensity of bands quantitated from 3-4 independent experiments, and the amounts of HA-DAT were normalized by TH. Asterisks indicate significant differences compared to Saline group, $p < 0.05$ (Student's unpaired t-test [n = 5]). *ns* indicates no significant difference.

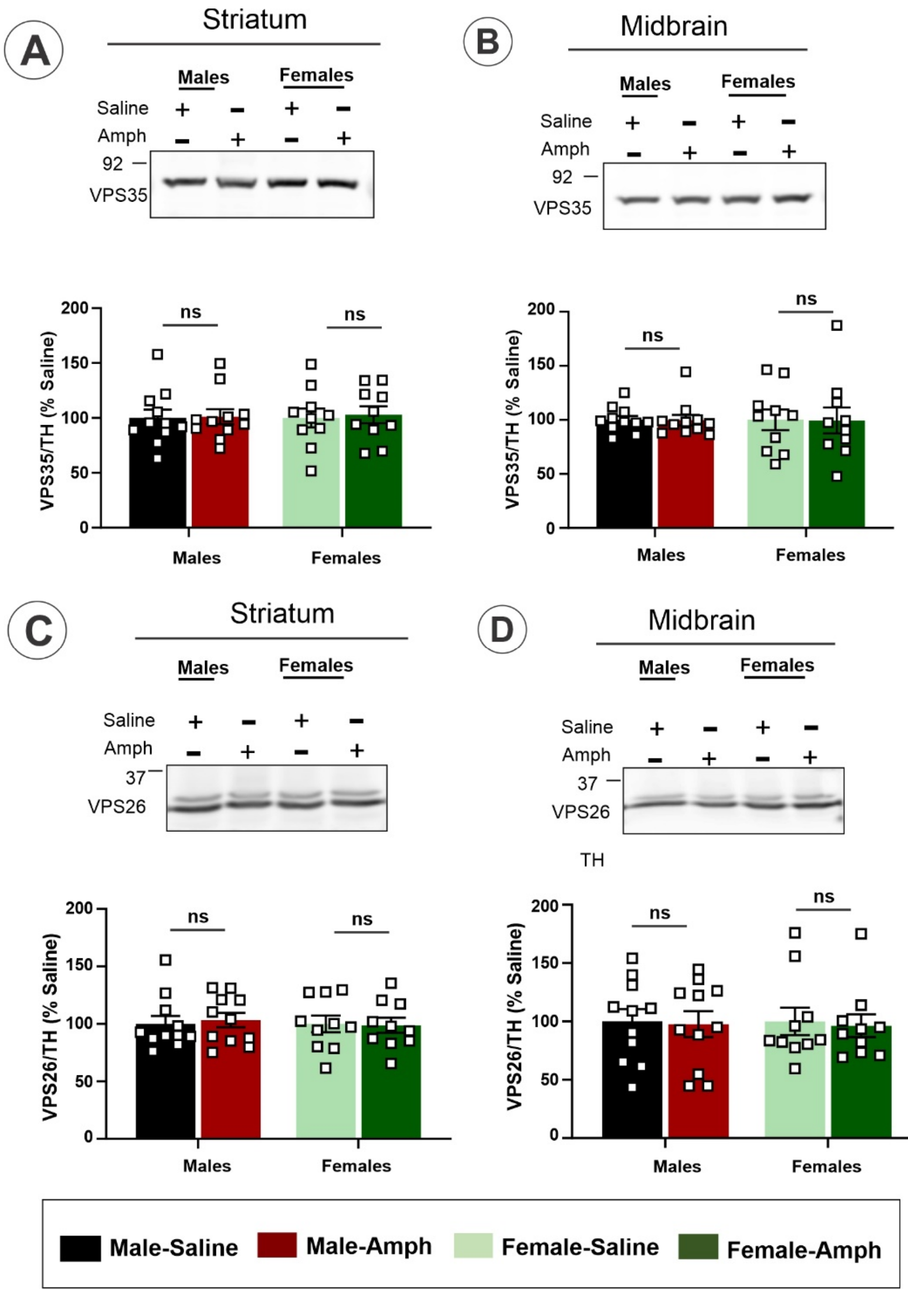


**Figure S3. Amph challenge does not affect the amounts of retromer complex in sensitized mice.**

Striatal synaptosomes (**A** and **C**) and MB tissue lysates (**B** and **D**) were electrophoresed and probed by immunoblotting with antibodies to VPS35 (**A-B**), VPS26 (**C-D**). Representative immunoblots are shown Bar graphs represent mean values (with SEMs) of VPS35/26 band intensities normalized to TH. Quantification of band intensities revealed no significant differences between male and female mice. Data are from three or more independent experiments. *ns, no significant difference.*

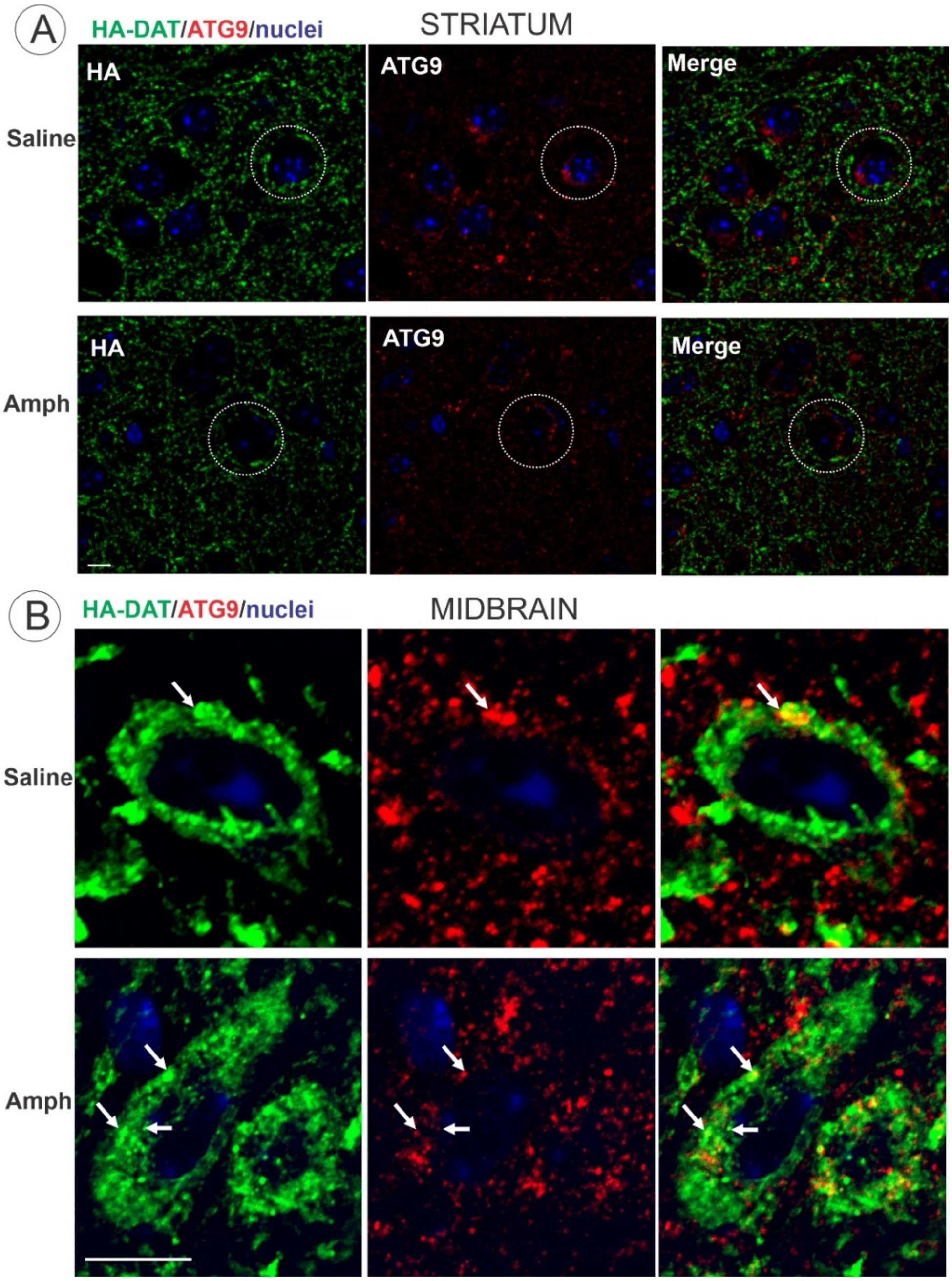


**Figure S4. Lack of the autophagic marker ATG9 in DA axons.**

Brains were fixed by cardiac perfusion with PFA and prepared for sectioning as described in Materials and Methods. Sagittal cryosections were co-labeled with rabbit or mouse HA11 antibodies and antibodies against ATG9 (**A-B**), followed by fluorophore-conjugated secondary antibodies. 3D stacks of confocal images were acquired through 640 nm (*red*, ATG9), 488 nm (*green,* HA-DAT), and 405 nm (*blue*, Hoechst) channels from striatum (**A**) and midbrain (**B**). White circles represent examples of characteristic perinuclear ATG9 labeling in HA-DAT-negative (nondopaminergic) cells. Arrow show examples of overlap of ATG9 and HA-DAT fluorescence in neuronal soma. All images are maximum intensity projections of three consecutive x-y confocal sections. Scale bars: **A**, 10 μm; **B**, 2 μm.
